# A changing hierarchy of environmental time cues guide development of the migratory pest moth *Loxostege sticticalis*

**DOI:** 10.64898/2025.12.26.696557

**Authors:** Changning Lv, Zun Xu, Yiyang Zhang, Hui Qiu, Jingxian Zhou, Fajun Chen, Xingfu Jiang, Abhishek Chatterjee, Guijun Wan

## Abstract

Insects use environmental cycling cues to align developmental progression. In holometabolous insects, the progression from larva to adulthood entails decision points, i.e., larval diapause and adult eclosion, that engage circadian timing mechanisms. What environmental time cues dynamically interact to regulate these developmental checkpoints lying at the intersection of the orthogonal timing systems, remains elusive. We used the seasonally migratory insect *Loxostege sticticalis* to disentangle the influence of photoperiod regimes and temperature conditions at distinct checkpoints of larval growth and metamorphosis. We here employed sensory conflict paradigms to reveal that the relative dominance of these cues’ changes over ontogeny. Our results confirm that diapause is induced primarily by the photoperiod during early larval development, rather than by the diel thermoperiod. In contrast, the circadian clock-controlled adult eclosion, normally locked to the first hour of the daybreak, is jointly controlled by light and temperature cycles. An increase in the constant temperature advanced the peak timing of daily eclosion. On the other hand, the interval timing mechanism regulating the duration of the eclosion event is tuned specifically by the night temperatures. Together, photoperiod dominates early developmental decisions, but thermoperiod gains weight at later stages. This shifting hierarchy provides a mechanistic framework for understanding phenological plasticity. Since the timing of eclosion presents a window of increased vulnerability for this pest insect, our results could help to optimize the daily timing of pest management intervention under variable thermal and light conditions.

## 1 Introduction

Earth’s revolution drives pronounced seasonal changes, whereas its rotation creates ∼24-h daily cycles of light and darkness, as well as temperature. Organisms adapt to variations in photoperiod and temperature fluctuations, enabling them to survive unfavorable seasonal conditions and anticipate optimal timings for their behaviors (Brady et al. 2021; Hidalgo and Chiu 2024). Light is the primary zeitgeber for circadian clocks, yet temperature ranks among the most potent abiotic modulators, directly altering clock pace and output (Saunders 2002; Beer and Helfrich-Förster 2020). Temperature cycles can entrain circadian clocks in complete absence of light cues (Lankinen and Riihimaa 1997; Glaser and Stanewsky 2005). The precise temporal coordination of biological processes by endogenous circadian clocks is essential for fitness and survival across taxa (Krittika and Yadav 2019; Patke et al. 2020).

In insects, evolutionarily conserved yet highly plastic molecular clocks orchestrate diverse physiological and behavioral processes (Patke et al. 2020). Diapause, a key adaptive strategy for surviving unfavorable conditions, manifests as hormonally controlled developmental arrest triggered by predictive environmental cues—chiefly photoperiod and temperature (Beck 1983; Schmidt and Conde 2006; Saunders 2020; Cui et al. 2025). Temperature profoundly modifies photoperiodic diapause induction (Saunders 1971; Yokoyama et al. 2021). Thermoperiods and naturally fluctuating temperature regimes have been reported to induce diapause more effectively than constant temperatures. Moreover, in species such as the European corn borer *Ostrinia nubilalis*, studies of combined daily light and temperature cycles suggest that the temperature during the cool dark phase may exert a greater influence on diapause regulation than the warm light phase (Fantinou and Kagkou 2000; Saunders 2013). Adult emergence (adult eclosion)—the final step of holometabolous metamorphosis—remains one of the most striking circadian-gated events in insects (Kalmus 1935; Bünning 1935; Saunders 2002). Eclosion is tightly restricted to specific times of day, producing rhythmic peaks that vary mainly with photoperiod, and thermoperiod (Pittendrigh 1954; Pittendrigh and Skopik 1970; Lankinen 1993; Numata and Tomioka 2023). In soil-pupating species such as the onion fly *Delia antiqua*, temperature cycles often dominate over light as the primary zeitgeber (Watari 2002, 2005; Tanaka and Watari 2003, 2011, 2025).

The beet webworm, *Loxostege sticticalis* (Lepidoptera: Crambidae), is a devastating migratory pest between 36° and 55° N, capable of massive outbreaks that inflict severe damage on crops and vegetables across Eurasia and North America (Zhang et al. 2008; Jiang et al. 2019). It overwinters as diapausing final-instar larvae in soil cocoons (Huang et al. 2009; Cui et al. 2025). Although extensive research has underpinned forecasting and control strategies for this species (Zhang and Jiang 2023), the chronobiology of its physiology and behavior remains largely unexplored. In particular, information on the adult eclosion rhythm of *L. sticticalis* remains limited.

Here, we investigated larval diapause induction in *L. sticticalis* under conditions in which seasonal information conveyed by photoperiodic and thermal cues is decoupled. We further characterized adult eclosion rhythms under constant and diel cycling temperature regimes, with and without light-dark cues. By elucidating the chronophysiological links between environmental signals and phenotypic timing, this study advances a mechanistic understanding of how seasonal cues are integrated across developmental stages in a migratory pest, and may help to inform phenological interpretation and forecasting under variable environmental conditions.

## 2 Materials and methods

### 2.1 Insect colony

Larvae of *L. sticticalis* were collected from the fields in Kangbao County, Hebei Province, China (41.85°N, 114.26°E) and transferred to the lab. The laboratory colony was established and maintained for over three generations on fresh leaves of *Chenopodium album* in climate chambers (RXZ-436; Ningbo Jiangnan Ltd., Ningbo, China) at 22 ± 0.4 °C, 70 ± 5% relative humidity and a photoperiod of 16 hrs light: 8 hrs dark (L16: D8; lights on at 7:00, off at 23:00; ∼1000 lux). Under these conditions, the complete life cycle of *L. sticticalis* lasted for around 48 days (Fig. 1a; Xu et al. 2025). Beet webworms were maintained in the climate chambers until further use.

**Fig. 1.**
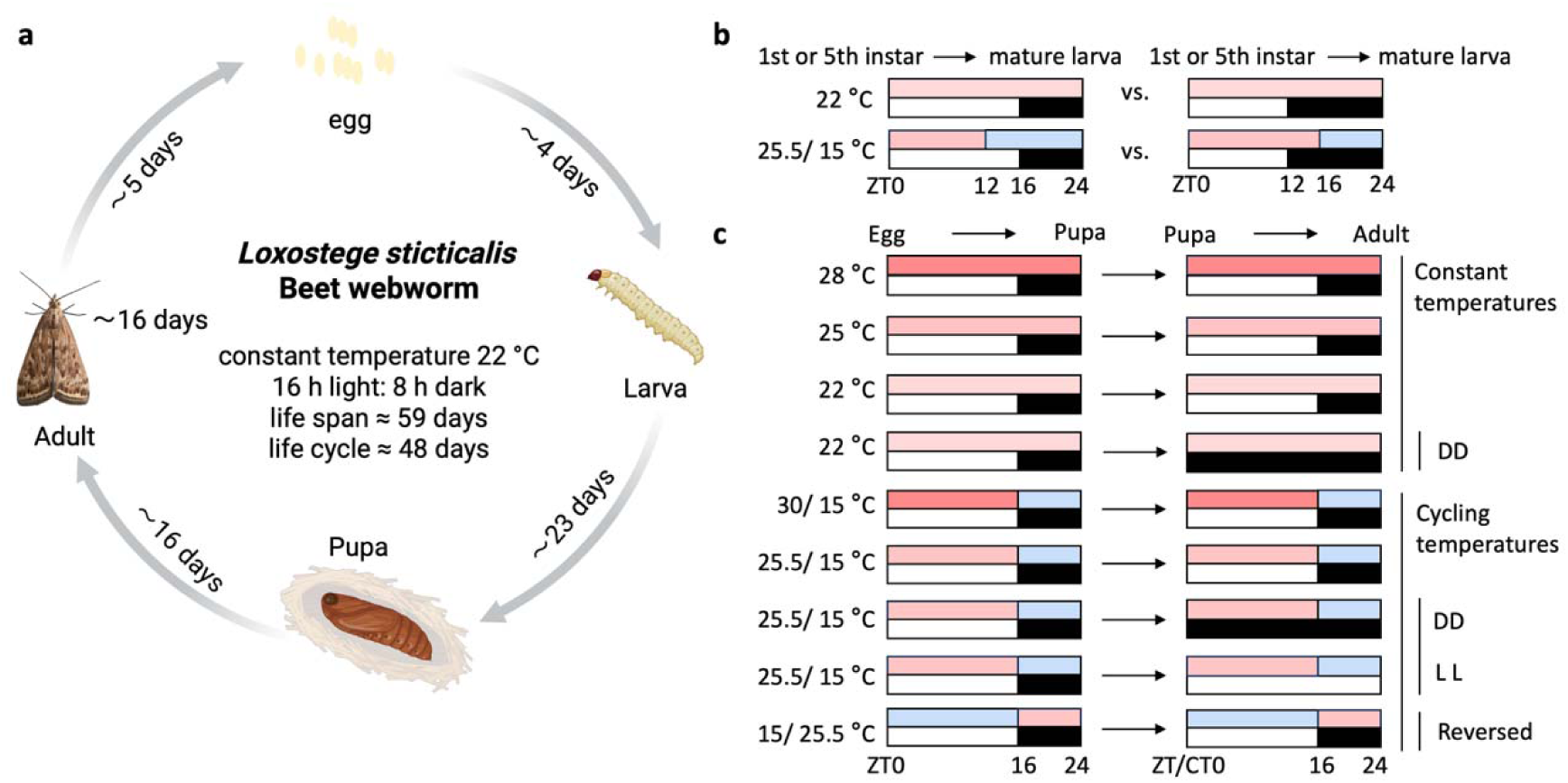
Life cycle of beet webworm *L. sticticalis* under constant temperature 22 °C and 16 hrs light: 8 hrs dark (data from Xu et al. 2025) (**a**). Temperature conditions and light regimes used in the diapause (**b**) and eclosion assay (**c**). “∼” and “≈” denote approximate values. Colorful bars represent temperature settings, with warmer/ cooler colors indicating higher or lower temperatures. White and black bars denote objective/ subjective day and night. ZT, zeitgeber time; CT, circadian time.

### 2.2 Diapause assay of *L. sticticalis* under different combination of photoperiods and temperatures

To induce diapause, first- and fifth-instar larvae reared at constant 22 °C under L16: D8 were transferred to 12 hrs light: 12 hrs dark (L12: D12; lights on at 7:00, off at 19:00; ∼1000 lux) while all other conditions remained unchanged. To examine the relative dominance of photoperiod versus thermoperiod in diapause induction, conflicting cues were applied. First- and fifth-instar larvae from the original constant 22 °C under L16: D8 colony were transferred to chambers providing either (i) L16: D8 combined with a 12 hrs 25.5 °C light/ 12 hrs 15 °C dark (warm phase started at 7:00, cool phase started at 19:00), or (ii) L12: D12 combined with 16 hrs 25.5 °C light/ 8 hrs 15 °C dark thermoperiod (warm phase started at 7:00, cool phase started at 23:00). The incidence of diapause in mature larvae was then recorded. Individuals whose prepupal developmental duration was several times longer than that of normally developing individuals at the same temperature were classified as diapausing individuals (Fig. 1b; Huang 2009).

### 2.3 Eclosion rhythm assay of *L. sticticalis* under different combination of light regimes and temperatures

To compare the effects of constant versus cycling temperatures on adult eclosion rhythm in *L. sticticalis* under L16: D8, we used three constant temperatures (28 °C, 25 °C, and 22 °C; Tang et al. 2016) and two thermoperiods 16 hrs 30 °C light/ 8 hrs 15 °C dark (30/ 15 °C; warm phase started at 7:00, cool phase started at 23:00; daily average temperature = 25 °C), mimicking May-September temperature conditions in Datong, Shanxi Province (40.08°N, 113.31°E), and 16 hrs 25.5 °C light/ 8 hrs 15 °C dark (25.5/ 15 °C; warm phase started at 7:00, cool phase started at 23:00; daily average temperature = 22 °C), mimicking the same month period in Qiqihar, Heilongjiang Province (47.36°N, 123.92°E) (Fig. 2). To determine whether the cycling temperature is sufficient to entrain the eclosion rhythm, larvae were reared under a 25.5/ 15 °C thermoperiod aligned with the L16: D8 photoperiod until pupation. And then newly formed pupae were gently transferred during either the subjective night or subjective day to identical chambers but set to constant darkness (DD) or constant light (LL) until eclosion. In addition, constant 22 °C under DD was used to assess the endogenous circadian rhythm and as a non-photic reference for the groups exposed to thermoperiodic cycles. Pupae at 13 days after pupation from the 22 °C colony (L16: D8) were transferred to chambers programmed for DD during subjective night. The average DD treatment duration was around 2.47 days. To assess the relative dominance of light and thermal cues, an additional cohort was reared under a reversed thermoperiod (15/ 25.5 °C; 16 hrs 15 °C light/ 8 hrs 25.5 °C dark; daily average temperature = 18.5 °C) (Fig. 1c).

**Fig. 2.**
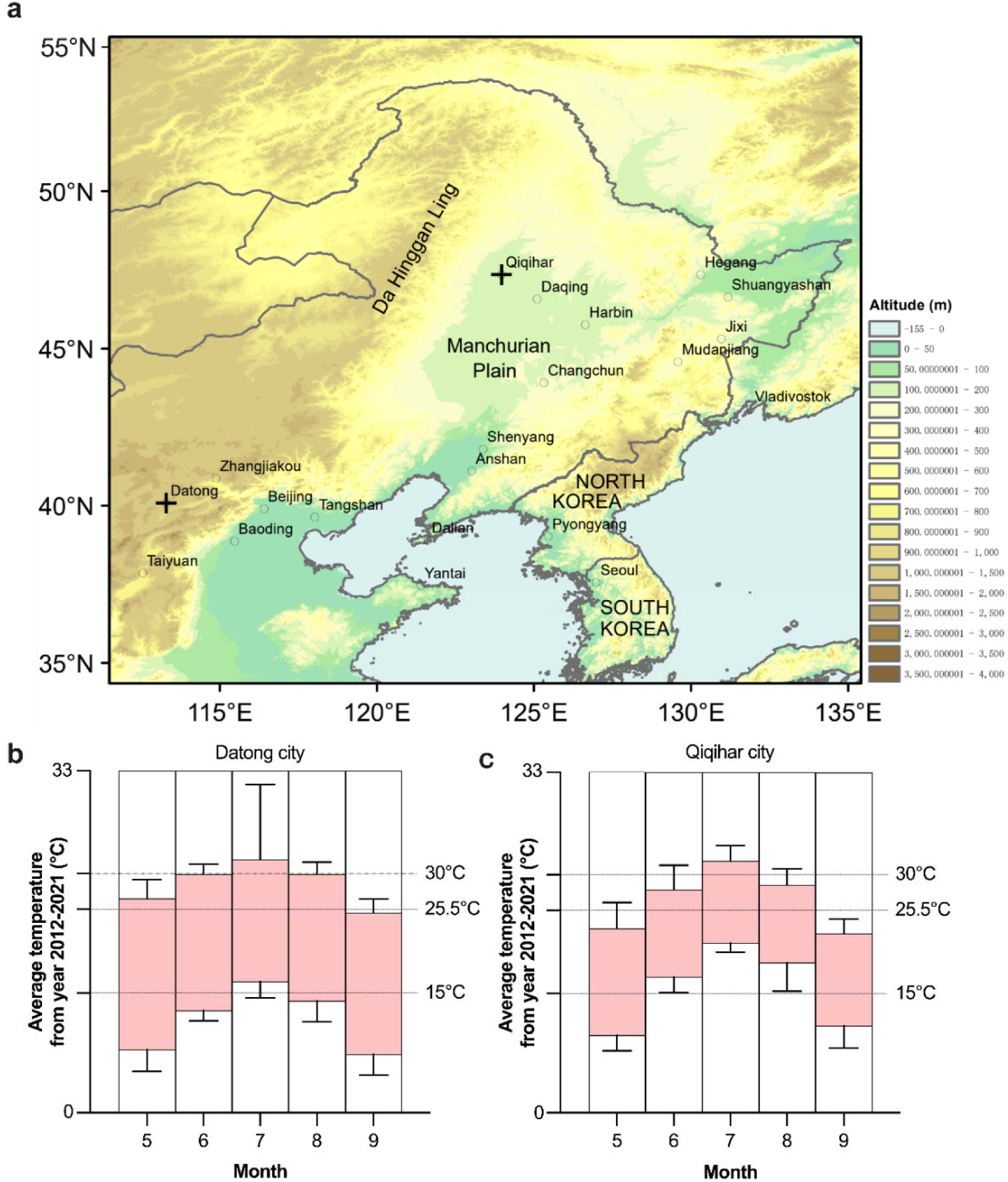
Regional map of parts of North China and Northeast China, and monthly temperature variations in cities with reported *Loxostege sticticalis* occurrences. **a** Cross symbols indicate the two cities, Datong City and Qiqihar City. The map includes both mountainous and plain areas, and was generated using ArcGIS. **b**, **c** Monthly average highest and lowest temperatures for Datong (**b**) and Qiqihar (**c**) from May to September, based on data from 2012 to 2021 (timeanddate.com). Pink shading indicates the temperature ranges associated with *L. sticticalis* outbreaks. Error bars represent the standard deviation (SD) of the highest and lowest temperatures. This visualization demonstrates that the selected temperature ranges are representative of conditions within the outbreak regions.

Pupal development of *L. sticticalis* was checked daily. Once pupal color changed from yellow to brown, individual pupae were transferred to ultra-clear glass tubes (diameter 2.4 cm, length 9.5 cm, wall thickness 0.2 cm) open at both ends and sealed with moisture cotton plugs. Each tube was uniquely numbered. Adult eclosion was continuously monitored using a custom infrared imaging system (Chinese patent ZL202120866706.1; Lv et al. 2025). Tubes containing pupae were placed in the monitoring device, and eclosion events were recorded with infrared cameras. Zeitgeber time ZT0 was defined as lights-on (07:00) and ZT16 as lights-off (23:00). Emerged adults were removed daily at 17:00 (ZT10), sexed by examination of abdominal terminalia, and their eclosion timing was determined from video recordings. Eclosion onset was defined as the moment the pupal case split and the adult head became visible, and completion as the point at which the adult had fully exited the pupa case. Eclosion duration was calculated as the interval between onset and completion.

### 2.4 Statistical analysis

Figures were generated using GraphPad Prism 10. Normality was assessed with the Shapiro-Wilk test (n ≤ 50). Diapause incidence was compared across treatments using χ² tests: Pearson χ² (asymptotic, two-sided) when the minimum expected count exceeded 5; continuity-corrected χ² when the minimum expected count was 1-5; and Fisher’s exact test (two-sided) when the minimum expected count was below 1. Differences in eclosion time distribution were evaluated using the exact two-sample Kolmogorov-Smirnov test and the Kruskal-Wallis test. Central tendencies of eclosion timing were compared across photoperiod-temperature treatments using median values with interquartile ranges, and differences among groups were assessed separately for females and males using Kruskal-Wallis tests with Bonferroni correction at adj *p* < 0.05 (SPSS 29.0). Eclosion duration between treatments was compared using One-way ANOVA and tested by post hoc Tukey HSD test at adj *p* < 0.05 (https://www.icalcu.com/stat/anova-tukey-hsd-calculator.html).

## 3 Results

### 3.1 Diapause incidence of *L. sticticalis* under different combination of photoperiods and temperatures

Under constant 22 °C and L16: D8, only 3.0 % of larvae treated from the 1st instar entered diapause, whereas none of the larvae treated at the 5th instar did so (Fig. 3a). In contrast, under constant 22 °C and L12: D12, 97.2 % of larvae treated from the 1st instar entered diapause, compared with only 6.1 % of those treated at the 5th instar (Fig. 3b). These results indicate that a short-day photoperiod (L12: D12) induces diapause in younger larvae far more effectively than in larvae first exposed at the 5th instar.

**Fig. 3.**
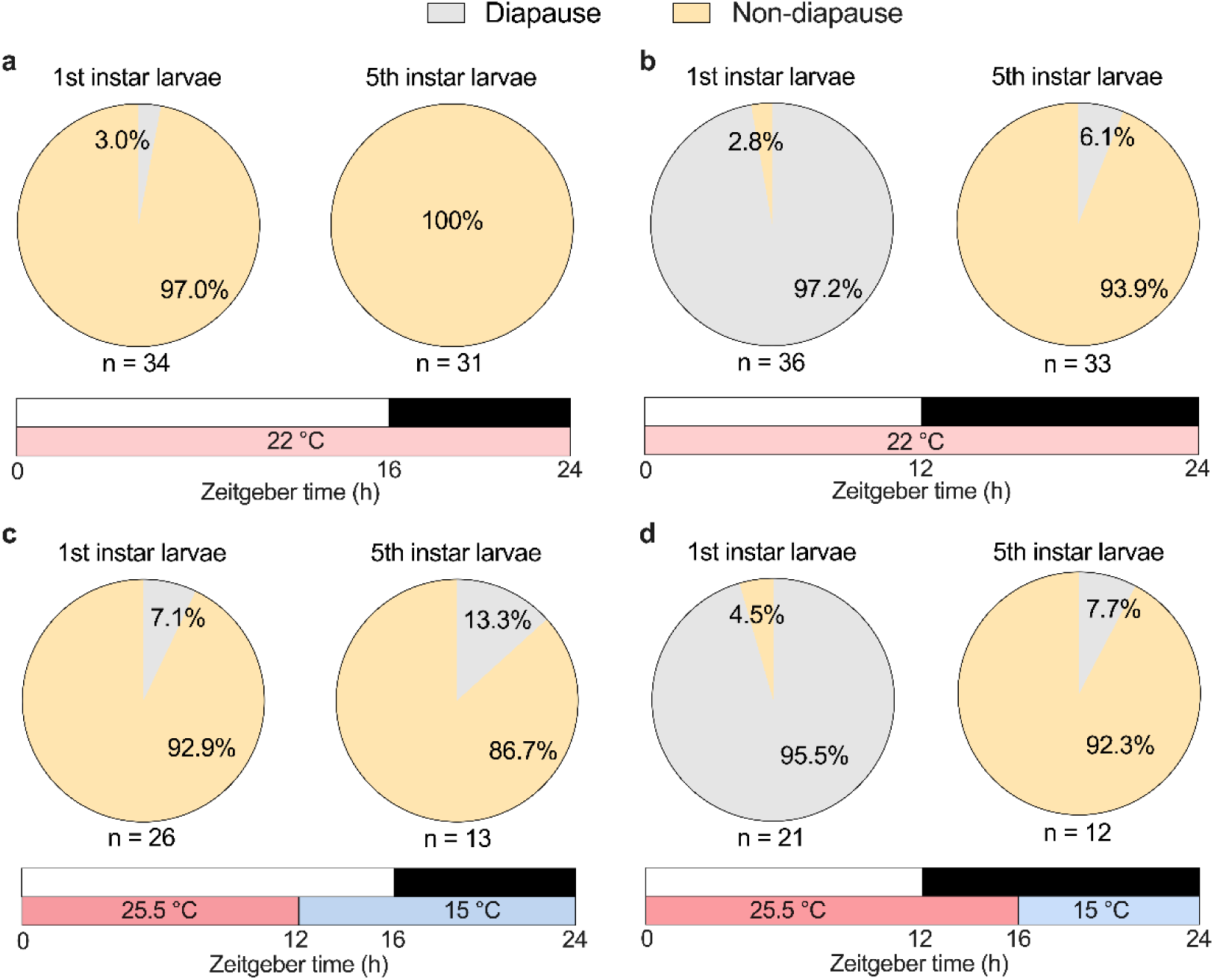
Diapause status of final-instar *L. sticticalis* larvae under different temperature and photoperiod combinations applied from the 1st or 5th instar. **a**, **b** Constant temperature (22 °C) under long-day (L16: D8; **a**) and short-day (L12:D12; **b**) photoperiods, with treatments applied from the 1st or 5th instar. **c** Cycling temperature regime (25.5/ 15 °C) combined with a long-day photoperiod (L16: D8): 16 hrs 25.5 °C light/ 8 hrs 15 °C dark. **d** Cycling temperature regime (25.5/ 15 °C) combined with a short-day photoperiod (L12: D12): 12 hrs 25.5 °C light/ 12 hrs 15 °C dark. Grey: diapause; yellow: non-diapause. White/ black bars: day/ night. Warmer/ cooler color bars: higher/ lower temperature. n: sample size.

Under the 12 hrs 25.5 °C light/12 hrs 15 °C dark thermoperiod with L16: D8, 7.1 % of larvae treated from the 1st instar entered diapause, compared with 13.3 % of those treated only at the 5th instar (Fig. 3c). Under the 16 hrs 25.5 °C light/8 hrs 15 °C dark thermoperiod with L12:D12, 95.5 % of larvae treated from the 1st instar entered diapause, whereas only 7.7 % of the 5th-instar-treated larvae did so (Fig. 3d). Chi-square tests showed that diapause incidence differed significantly between larvae treated from the 1st instar and those treated only at the 5th instar under short-day conditions (L12: D12; *p* < 0.001), but not under long-day conditions (L16: D8; *p* ≥ 0.852) (Fig. 3; Table 1). Across both constant and cycling temperatures, diapause incidence in larvae treated from the 1st instar (but not in those treated only at the 5th instar, *p* ≥ 0.493) differed significantly between long-day and short-day photoperiods (*p* < 0.001; Fig. 3; Table 2). In contrast, diapause incidence did not differ between constant and cycling temperature regimes under any photoperiod or instar treatment (*p* ≥ 0.082; Fig. 3; Table 3). Together, these results demonstrated that photoperiod, rather than cycling temperature, was the primary cue inducing diapause in younger larvae of *L. sticticalis*.

**Table 1.** Chi-square test of diapause incidence in *L. sticticalis* larvae treated from different larval stages (1st vs. 5th instar) under identical photoperiod-temperature conditions.

| Temperature | L16: D8 (*p* value) | L12: D12 (*p* value) |
| --- | --- | --- |
| Constant temperature: 22 °C | 1.00 | <0.001*** |
| Cycling temperature: 12 hrs 25.5 °C light/ 12 hrs 15 °C dark or 16 hrs 25.5 °C light/ 8 hrs 15 °C dark | 0.852 | <0.001*** |

**Note:** Pearson's Chi-square test (asymptotic, two-sided) was used when the minimum expected count was >5; the continuity-corrected Chi-square test (asymptotic, two-sided) was used for expected counts between 1 and 5; and Fisher's exact test (two-sided, exact) was applied when any expected count was <1. \*\*\**p* < 0.001.

**Table 2.** Chi-square test of diapause incidence in *L. sticticalis* larvae exposed to different photoperiods (L16: D8 vs. L12: D12) from the 1st or 5th instar under identical temperature conditions.

| Temperature | Treated from the 1st instar (*p* value) | Treated from the 5th instar (*p* value) |
| --- | --- | --- |
| Constant temperature: 22 °C | <0.001 *** | 0.493 |
| Cycling temperature: 12 hrs 25.5 °C light/ 12 hrs 15 °C dark or 16 hrs 25.5 °C light/ 8 hrs 15 °C dark | <0.001 *** | 0.792 |

**Note:** Pearson's Chi-square test (asymptotic, two-sided) was used when the minimum expected count was >5; the continuity-corrected Chi-square test (asymptotic, two-sided) was used for expected counts between 1 and 5; and Fisher's exact test (two-sided, exact) was applied when any expected count was <1. \*\*\**p* < 0.001.

**Table 3.** Chi-square test of diapause incidence in *L. sticticalis* larvae exposed to different temperature conditions (constant 22 °C vs. cycling 12 hrs 25.5 °C light/ 12 hrs 15 °C dark or 16 hrs 25.5 °C light/ 8 hrs 15 °C dark) from the 1st or 5th instar under identical photoperiod conditions

| Photoperiod | Treated from the 1st instar (*p* value) | Treated from the 5th instar (*p* value) |
| --- | --- | --- |
| L16: D8 | 0.811 | 0.082 |
| L12: D12 | 1.00 | 0.126 |

**Note:** Pearson's Chi-square test (asymptotic, two-sided) was used when the minimum expected count was >5; the continuity-corrected Chi-square test (asymptotic, two-sided) was used for expected counts between 1 and 5; and Fisher's exact test (two-sided, exact) was applied when any expected count was <1.

### 3.2 Eclosion rhythms of *L. sticticalis* under constant temperature conditions

Under the three constant temperature treatments with a L16: D8 photoperiod, both female and male *L. sticticalis* adults displayed diel eclosion rhythms with two peaks. At 28 °C, the major peak occurred at ZT17-ZT18 in both sexes (females: 22.6 %; males: 35.1 %; Fig. 4a, b). The secondary peak was observed at ZT21-ZT22 in females (19.4 %) and at ZT18-ZT19 in males (18.9 %). Across the 24 h cycle, only 9.7 % of female eclosion and 5.4 % of male eclosion occurred during the light phase, with the remainder in the dark phase (Fig. 4a, b). Both peaks in females and males occurred within the dark phase.

**Fig. 4.**
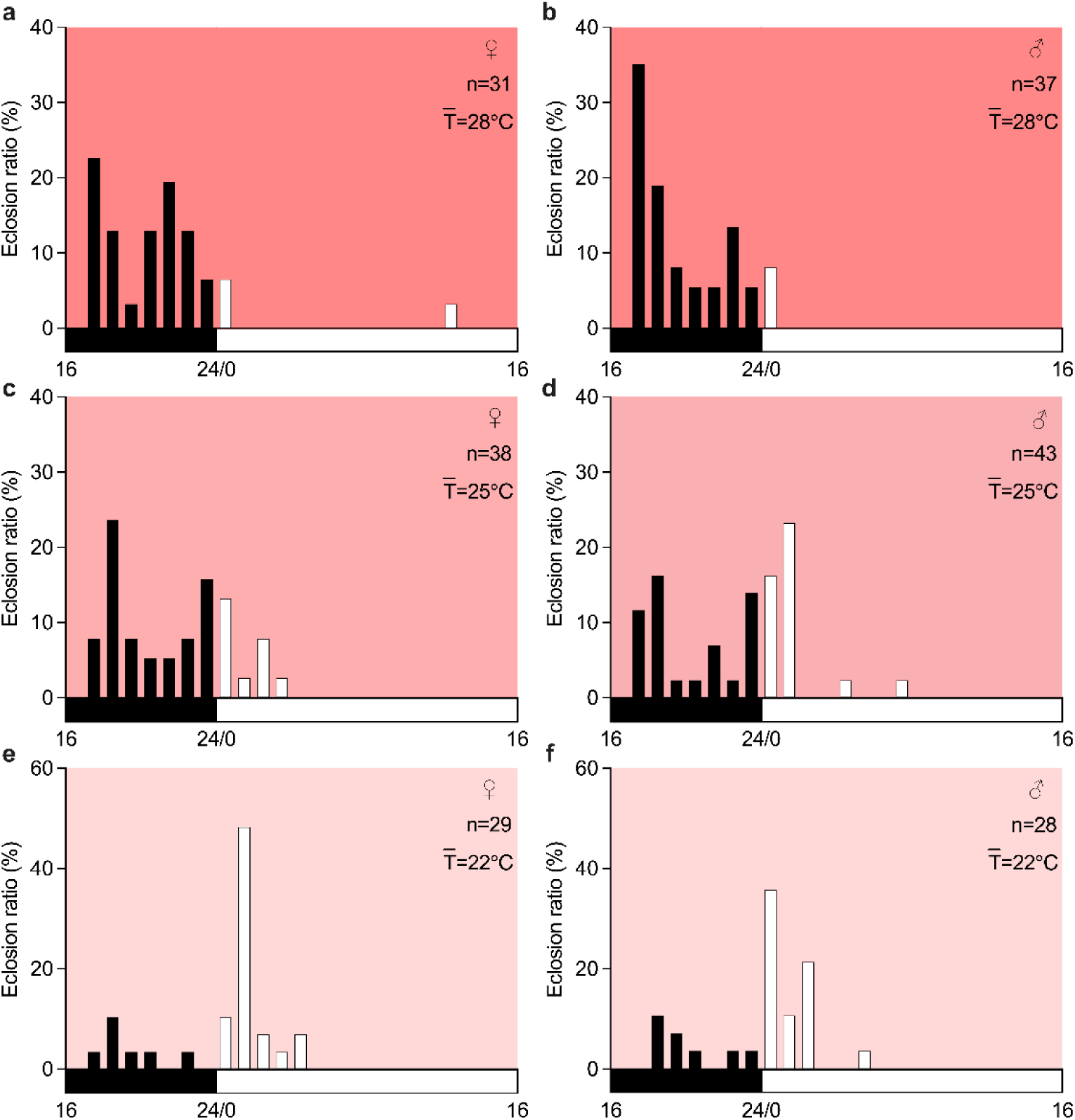
Eclosion rhythms of *L. sticticalis* under different constant temperature conditions with a 16 hrs light: 8 hrs dark photoperiod. **a, b** Female (**a**) and male (**b**) adults at 28 °C. **c, d** Female (**c**) and male (**d**) adults at 25 °C. **e, f** Female (**e**) and male (**f**) adults at 22 °C. Each column represents the hourly eclosion ratio within a 24 h cycle. ♀, female; ♂, male; n, sample size; TLJ, average temperature. Warmer color background: higher temperature. Horizontal bars indicate the objective day (white) and night (black).

At 25 °C, females showed a major peak at ZT18-ZT19 (23.7 %) and a secondary peak at ZT23-ZT24 (15.8 %), both in the dark phase (Fig. 4c). In males, the major peak occurred at ZT2-ZT3 (23.3 %) in the light phase, and secondary peaks appeared at ZT0-ZT1 and ZT18-ZT19 (16.3 %; Fig. 4d). Across 24 h cycle, 26.3 % of female eclosion and 44.2 % of male eclosion occurred in the light phase. The timing of the major peaks differed between sexes, both sexes showed a higher proportion of eclosion during the dark phase (Fig. 4c, d).

At 22 °C, the primary peaks for females and males occurred at ZT1-ZT2 and ZT0-ZT1, respectively (females: 48.3 %; males: 35.7 %; Fig. 4e, f), coinciding with the dark-light transition. Secondary peaks were not clearly defined. Across the cycle, 75.9 % of female eclosion and 71.4 % of male eclosion occurred in the light phase (Fig. 4e, f). In both sexes, the major peak occurred during the light phase.

Under DD at 22 °C, *L. sticticalis* adults continued to exhibit a discernible eclosion peak (average duration in DD: ∼2.47 d for both sexes). The major peak for females occurred at CT2-CT3 (25.0 %) and for males at CT3-CT4 (18.2 %; Fig. 5). Compared with the sharper peaks under LD conditions, eclosion timing in DD was more broadly distributed. The eclosion peaks in DD occurred later than those under LD, suggesting that the underlying circadian period may be slightly longer than 24 h under these conditions.

**Fig. 5.**
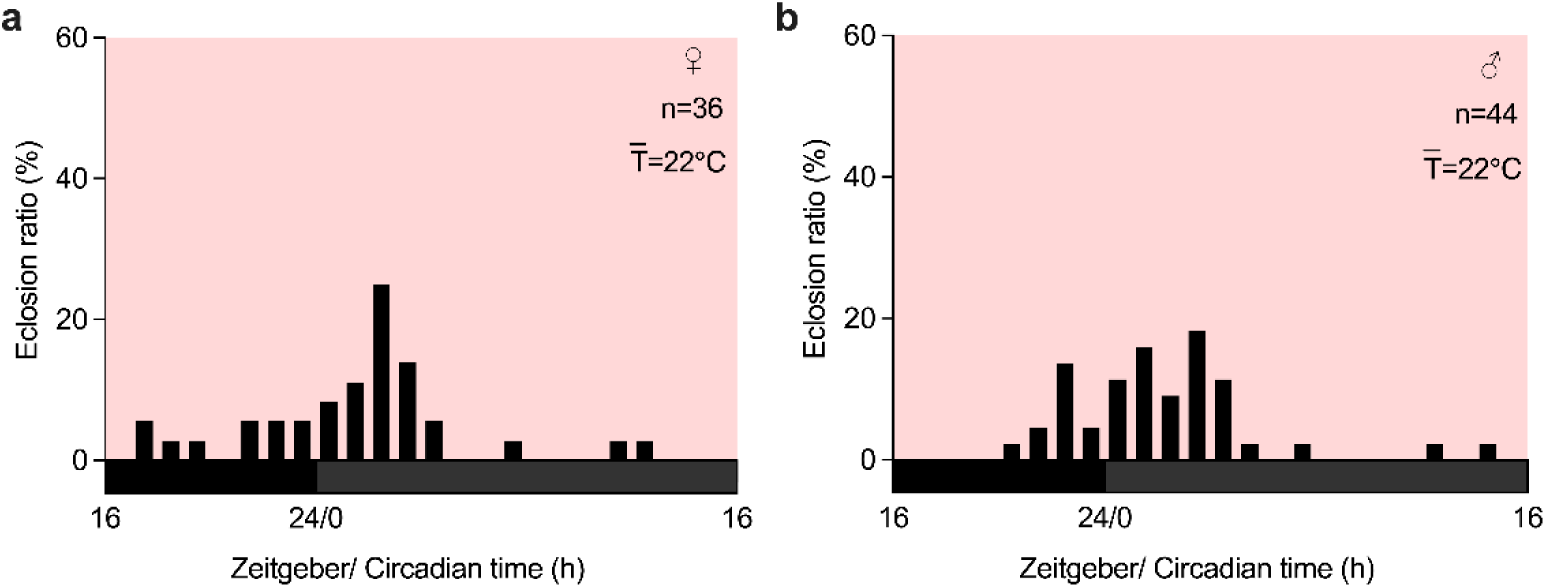
Eclosion rhythms of female (**a**) and male (**b**) *L. sticticalis* under constant darkness at 22 °C. Each column represents the hourly eclosion ratio within a 24 h cycle. ♀, female; ♂, male; n, sample size; TLJ, average temperature. Pink background: 22 °C. Horizontal bars indicate the subjective day (dark grey) and night (black).

### 3.3 Eclosion rhythms of *L. sticticalis* under cycling temperature with different light regimes

Under the two cycling temperature treatments (30/ 15 °C and 25.5/ 15 °C) with an L16: D10 photoperiod, both female and male *L. sticticalis* adults showed circadian eclosion rhythms with a single daily peak (Fig. 6a-d). Under 30/ 15 °C, the peak occurred at ZT0-ZT1 in both sexes (females: 70.0 %; males: 56.7 %), within the first hour of the light phase. Across the 24 h cycle, 76.7% of female eclosion and 60.0% of male eclosion occurred during the light phase (Fig. 6a, b).

**Fig. 6.**
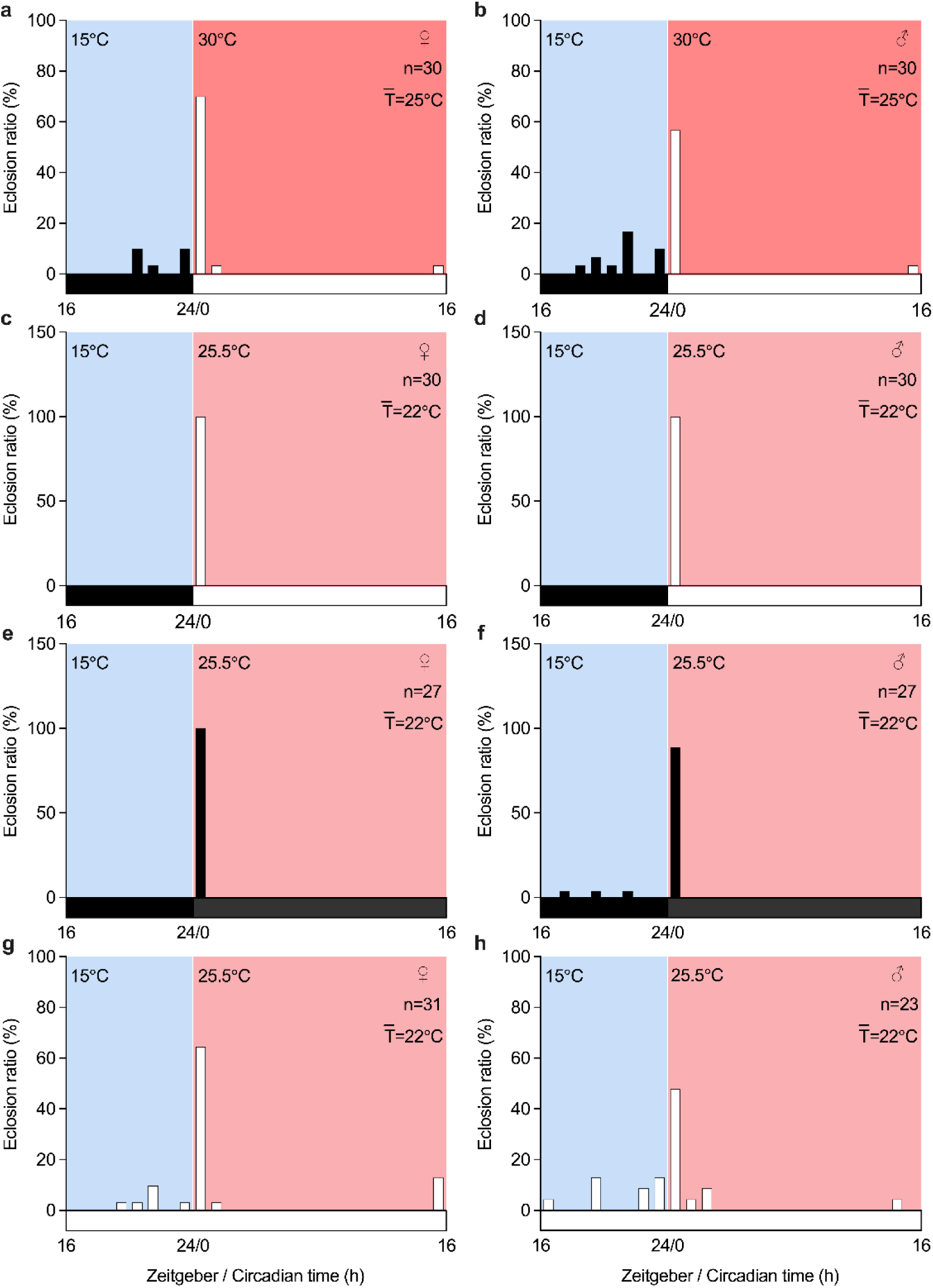
Eclosion rhythms of *L. sticticalis* under cycling temperatures combined with different light regimes. **a**, **b** Female (**a**) and male (**b**) adults under a cycling temperature regime of 30/ 15 °C combined with a long-day photoperiod (L16: D8). **c**, **d** Female (**c**) and male (**d**) adults under a cycling temperature regime of 25.5/ 15 °C combined with a long-day photoperiod (L16: D8). **e**, **f** Female (**e**) and male (**f**) adults under a cycling temperature regime of 25.5/ 15 °C under constant darkness. **g**, **h** Female (**g**) and male (**h**) adults under a cycling temperature regime of 25.5/ 15 °C under constant light. Each column represents the hourly eclosion ratio over 24 h. ♀, female; ♂, male; n, sample size; TLJ, average temperature. Warmer/ cooler color background: higher/ lower temperature. Horizontal bars indicate objective day (white)/ night (black), subjective day (dark grey)/ night (black), and constant light (white).

Under 25.5/ 15 °C, females and males also showed a single peak at ZT0-ZT1 (100 % in both sexes; Fig. 6c, d). At the same cycling temperature but under DD, the peak again occurred at ZT0-ZT1 (females: 100 %; males: 88.9 %; Fig. 6e, f). Under LL, the peak remained at ZT0-ZT1 (females: 64.5%; males: 47.8%; Fig. 6g, h).

To assess the relative influence of photoperiod and thermoperiod, we reversed their natural alignment by imposing a thermoperiod of 15 °C during the 16-h light phase and 25.5 °C during the 8-h dark phase while maintaining the L16: D8 photoperiod. Under this conflicting regime, both sexes eclosed mainly during the light phase (females: 85.0 %; males: 100 %), with peak eclosion at ZT5-ZT6 (females: 20.0%; males: 41.2%) (Fig. 7). These results indicate that eclosion remained closely aligned with the light-dark cycle despite the inverted temperature cycle.

**Fig. 7.**
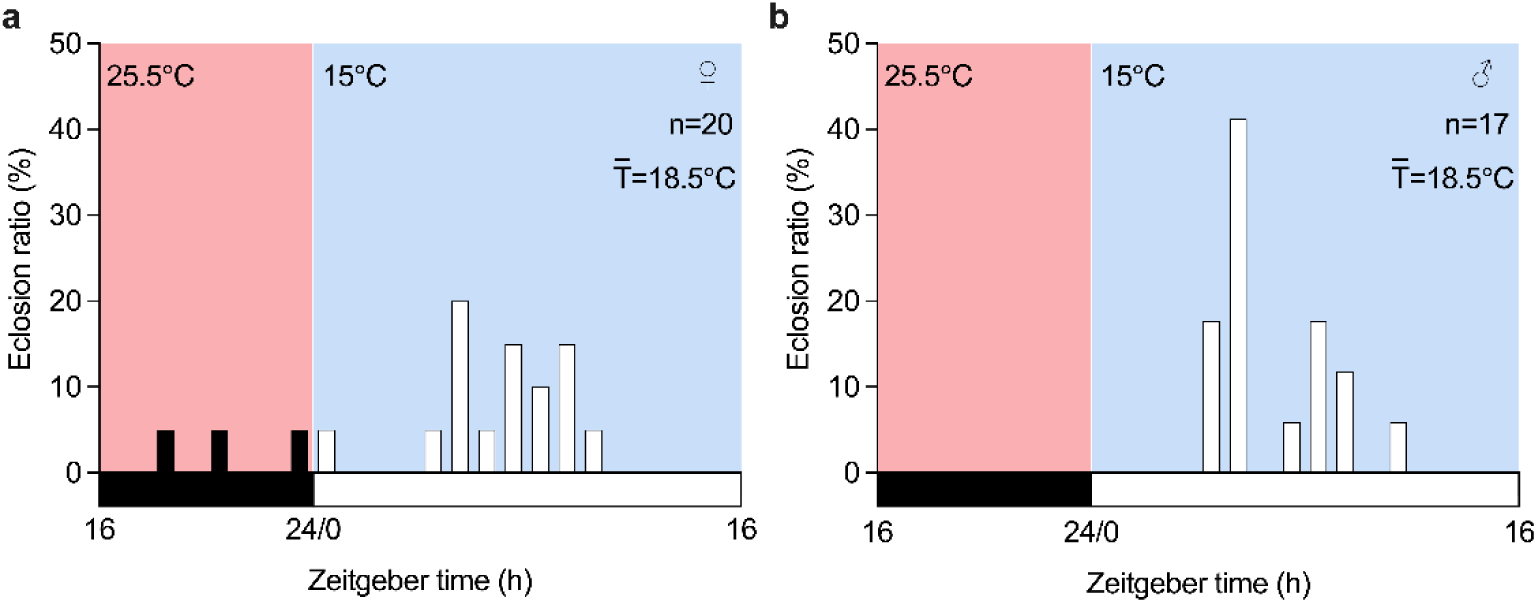
Eclosion rhythms of female (**a**) and male (**b**) *L. sticticalis* under a cycling temperature regime (16 hrs 15 °C light/ 8 hrs 25.5 °C dark), in which the temperature cycle was inverted relative to the light-dark cycle, resulting in higher temperatures at night and lower temperatures during the day. ♀, female; ♂, male; n, sample size; TLJ, average temperature. Warmer/ cooler color background: higher/ lower temperature. Horizontal bars indicate objective day (white) and night (black).

### 3.4 Eclosion time distribution of *L. sticticalis* under different light regimes and temperature conditions

To gain a more detailed understanding of the overall distributions of adult eclosion of *L. sticticalis*, the exact two sample K-S test was applied to examine differences in eclosion distribution between treatments and sexes. Under constant temperatures, eclosion distribution at 28 °C differed significantly from those at 25°C and 22 °C in females (*p* < 0.001), whereas no difference was detected between 25 °C and 22 °C (*p* = 0.161). In males, eclosion distributions at 28 °C and 25 °C both differed significantly from 22 °C (*p* < 0.001), while 28 °C and 25 °C did not differ significantly (*p* = 0.200).

Under cycling temperatures, significant difference was observed between 25.5/ 15 °C and 30/ 15 °C in females (*p* = 0.016) but not in males (*p* = 0.388). Compared with constant 22 °C, the 25.5/ 15 °C thermoperiod (mean 22 °C) differed significantly in females and males (*p* < 0.0497). Compared with constant 25 °C, the 30/ 15 °C thermoperiod (mean 25 °C) did not differ significantly in females (*p* = 0.236) but in males (*p* < 0.001). Under 25.5/ 15 °C with different light regimes (L16: D8, LL, DD), no significant distributional differences were detected (*p* > 0.489). Reversing the alignment between thermoperiod and photoperiod (25.5/ 15 °C vs. 15/ 25.5 °C) resulted in a significant difference in eclosion distributions (*p* < 0.001).

Given that the Kruskal-Wallis test captures differences in central tendency, it was additionally used to assess variation in median eclosion timing. For constant temperatures, females showed significant differences between 22 °C and both 28 °C and 25 °C (adj *p* < 0.001), while 28 °C and 25 °C did not differ (adj *p* = 1.000). Males showed significant differences between 28 °C and both 25 °C and 22 °C (adj *p* ≤ 0.002), while 25 °C and 22 °C did not differ (adj *p* = 1.000). For cycling temperatures, the difference between 25.5/ 15 °C and 30/ 15 °C was not significant in females and males (adj *p* = 1.000). Compared with 22 °C, the 25.5/ 15 °C treatment did not differ significantly for females and males (*p* = 1.000). Compared with 25 °C, the 30/ 15 °C treatment differed significantly for females and males (adj *p* ≥ 0.270). Under 25.5/ 15 °C with L16: D8, LL, and DD, no significant differences were detected (adj *p* = 1.000). Reversing thermoperiod-photoperiod alignment significantly altered median eclosion timing in both sexes (adj *p* ≤ 0.045) except with constant 22 °C under light-dark cycles and constant darkness in females (adj *p* ≥ 0.723) (Fig. 8).

**Fig. 8.**
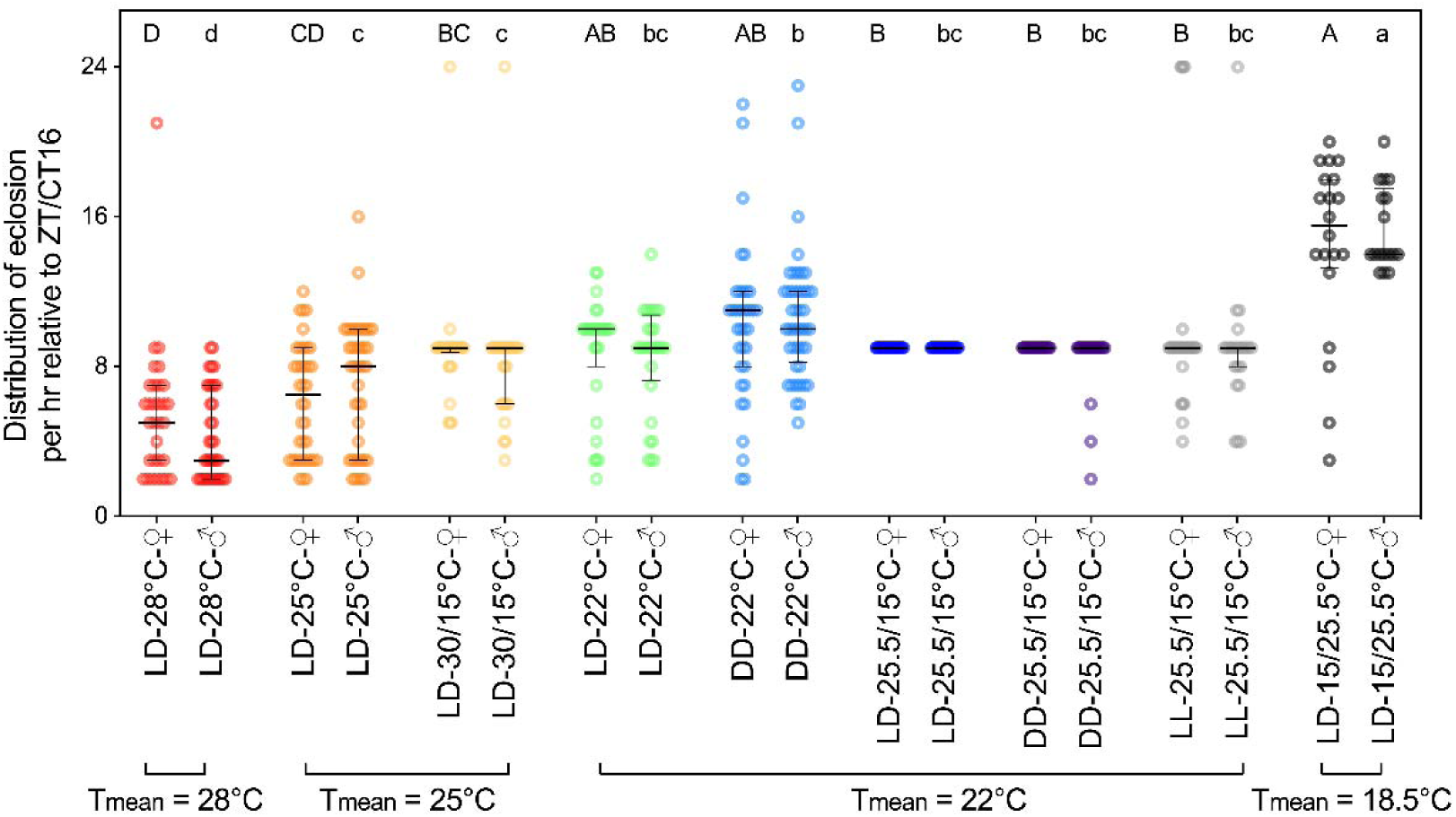
Central tendency comparison of eclosion rhythms in *L. sticticalis* under combination of different light regimes and temperature conditions. Solid lines represent median values, and error bars indicate interquartile ranges. ♀, female; ♂, male. U-shaped connectors mark groups with the same average temperature. Uppercase and lowercase letters indicate significant differences within females and within males, respectively, based on the Kruskal-Wallis test with Bonferroni correction at adj *p* < 0.05. On the x-axis, LD: 16 hrs light: 8 hrs dark; LL: constant light; DD: constant darkness. The numbers separated by “/” represent temperature cycles, and the temperature cycle time ratio is 16 hrs: 8 hrs.

### 3.5 Eclosion durations of *L. sticticalis* under different temperature treatments

The average eclosion duration of female and male adults was 1.70 and 1.73 min at 28 °C, 2.67 and 2.63 min at 25 °C, and 3.24 and 3.23 min at 22 °C, respectively. Under constant temperature, eclosion duration increased significantly as temperature decreased (adj *p* < 0.05). Under the cycling temperatures, females and males showed mean durations of 3.77 and 3.83 min at 30/ 15 °C and 3.67 and 3.66 min at 25.5/ 15 °C, respectively (Fig. 8). No significant difference was detected between the two thermoperiods (adj *p* > 0.05). When the daily mean temperature was matched (25 °C vs. 30/ 15 °C), eclosion duration of *L. sticticalis* was significantly longer under cycling than constant temperatures, but not in 22 °C vs. 25.5/ 15 °C group (Fig. 9).

**Fig. 9.**
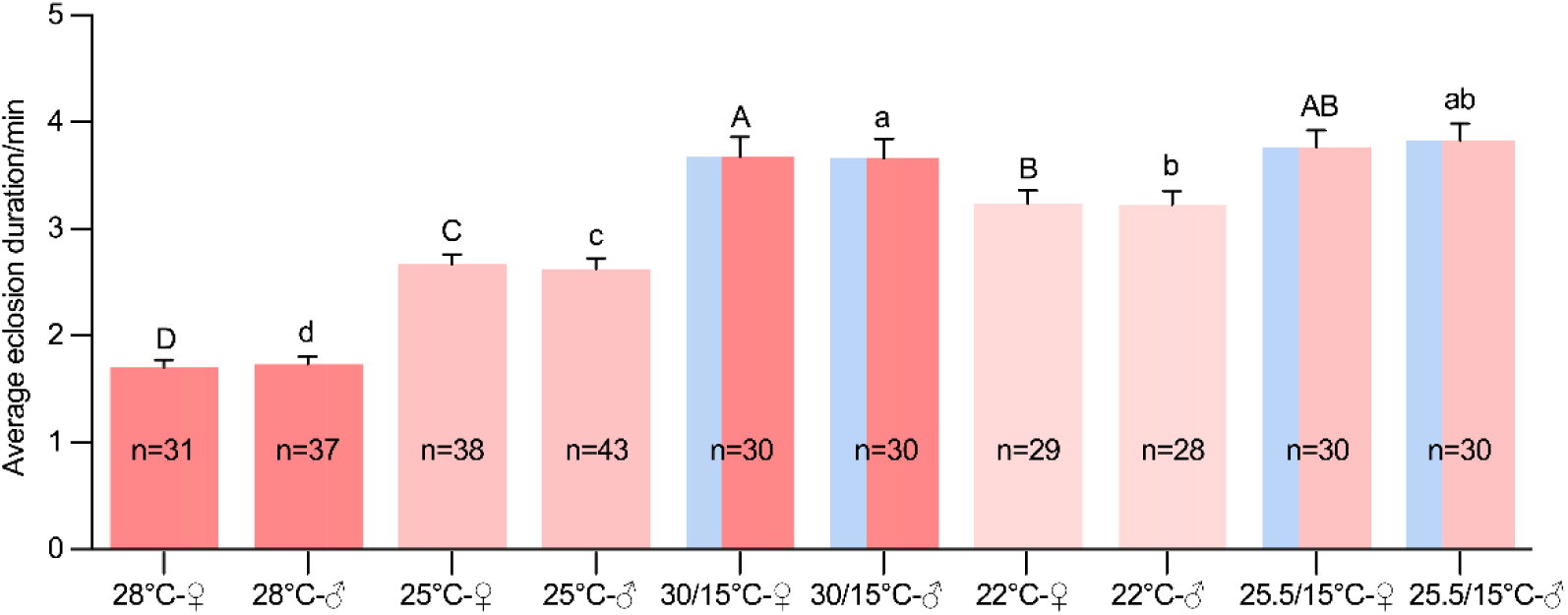
Average eclosion durations of *L. sticticalis* under constant and cycling temperature conditions with a 16 hrs light: 8 hrs dark photoperiod. Warmer/ cooler color bars: higher/ lower temperature. Error bars represent standard error (SE). Different uppercase and lowercase letters indicate significant differences among female and male treatments as determined by Tukey HSD test at adj *p* < 0.05. The numbers separated by “/” represent temperature cycles, and the temperature cycle time ratio is 16 hrs: 8 hrs. ♀, female; ♂, male; n, sample size.

## 4 Discussion

Accurate time perception is fundamental to insect survival, as endogenous biological clocks integrate environmental cues to align development and behavior with favorable temporal windows. Here, we investigated the migratory pest *Loxostege sticticalis*, which is notorious for outbreaks widely distributed in temperate regions worldwide, to compare the relative roles of photoperiodic and thermal cues in regulating larval diapause induction. We further characterized diel and circadian adult eclosion rhythms under a range of light regimes and temperature conditions, using eclosion as a classical output of the circadian system. Our results show that the relative influence of light and temperature cues changes across development, providing insight into how seasonal and daily timing mechanisms are integrated within a single life cycle. This stage-dependent shift in cue influence helps to explain the distinct timing patterns observed in *L. sticticalis* and provides a framework for interpreting its phenological responses under variable environmental conditions.

In migratory insect, adults often embark on flights the day after eclosion to locate more favorable habitats (Liu et al. 2021), while diapause represents an alternative strategy to withstand adverse conditions (Numata and Shintani 2023). Insect diapause is a fascinating phenomenon in insect chronobiology, and it is closely related to light and temperature. Goryshin et al. (1980) proposed that diapause induction in *L. sticticalis* larvae is primarily regulated by photoperiod, independent of temperature and latitude, and represents a classic long-day response type. In contrast, Saulich et al. (1983) investigated populations from two climatic zones in the former Soviet Union and demonstrated clear intraspecific variation: larvae from the steppe zone exhibited typical photoperiodic regulation of diapause, whereas populations from the forest-steppe zone entered early diapause irrespective of photoperiod. Using populations of *L. sticticalis* collected in China, Cui et al (2025) showed that under short-day condition, first-instar larvae show higher diapause incidence than fifth-instar larvae, indicating that early instars may represent a sensitive period for perceiving short-day cues associated with diapause induction. We observed a consistent pattern in our experiments. Under the same photoperiod, the introduction of diel temperature cycles did not significantly alter diapause incidence, indicating that photoperiodic cues may play a more prominent role than thermoperiodic cues in diapause induction in this species. Alternatively, it is also possible that the amplitude or minimum temperature of the imposed temperature cycles did not exceed the threshold required to modulate diapause responses (Fantinou and Kagkou 2000; Saunders, 2013).

Diel adult eclosion rhythms vary across insect species and can be broadly classified as unimodal or bimodal. A unimodal diel pattern has been reported in several insects, including *Plodia interpunctella*, *Spodoptera frugiperda* and *Nilaparvata lugens* (Kikukawa et al. 2013; Lv et al. 2025; Xuanyuan et al. 2025). In contrast, a bimodal diel eclosion rhythm has been described in the silkworm *Bombyx mori*, in which one of the daily peaks has been attributed to a direct response to light, consistent with a masking effect rather than endogenous circadian control (Shimizu and Matsui 1983). In *L. sticticalis*, we observed a similar bimodal diel pattern under constant temperature conditions. One eclosion peak consistently occurred within approximately three hours after lights-off, but this peak disappeared under constant darkness, indicating that it is unlikely to be generated by the endogenous circadian clock. Instead, its timing and light dependence are consistent with a masking effect induced by the light-dark transition (Rietveld et al. 1993; Lamont and Amir 2009; Gall and Shuboni-Mulligan 2022). The phase of this peak remained relatively stable across temperatures, whereas its amplitude declined at lower temperatures. By contrast, the second peak exhibited pronounced temperature sensitivity, with its phase progressively delayed as temperature decreased. When exposed to cycling temperature regimes, *L. sticticalis* displayed a unimodal and highly synchronized population-level eclosion rhythm. This shift from bimodal to unimodal diel expression underscores the strong synchronizing influence of thermoperiodic cues on eclosion timing. Notably, robust synchrony emerged only when both photoperiodic and thermoperiodic cues were present, consistent with the joint action of circadian gating and environmentally induced masking. In this respect, *L. sticticalis* differs from the Asian gypsy moth *Lymantria dispar*, which retains a bimodal diel eclosion rhythm under natural temperature cycles but becomes unimodal under laboratory constant temperatures (Ma et al. 1982; Cardé et al. 1996). Together, these comparisons indicate that the expression of unimodal or bimodal diel eclosion rhythms depends on how circadian control and environmental modulation interact across species and environmental contexts.

Cycling temperature regimes exerted a strong synchronizing influence on adult eclosion in *L. sticticalis*, even when light-dark cues were removed, consistent with thermoperiodic entrainment of population-level timing. This sensitivity to temperature cycles may be relevant to the ecology of *L. sticticalis*, which is most active in mid- to high-latitude inland regions where diel temperature amplitudes can be substantial (Hoikkala and Poikela 2022). Moreover, consistent with earlier work showing pronounced temperature dependence of survival and performance in this species (Cheng et al. 2015; Tang et al. 2016), we observed marked shifts in eclosion timing across thermal conditions. Under constant temperatures, increasing temperature shifted the major eclosion peak from the early morning towards late night, whereas under thermoperiods the population expressed a tightly clustered unimodal peak. Notably, synchrony was reduced under the warmer thermoperiod (30/15 °C) relative to 25.5/15 °C, which may reflect increased individual variation in clock output or in responsiveness to thermal cues under less favorable daytime temperatures (Cheng et al. 2015; Tang et al. 2016; Xu et al. 2025). When photoperiod and thermoperiod were experimentally misaligned, eclosion remained primarily aligned with the light-dark cycle and occurred predominantly during the photophase, even though daytime temperature was lower. This pattern is consistent with photoperiodic cues providing a robust reference for eclosion timing when environmental time cues conflict. Across treatments, we observed a temperature-dependent phase shift in eclosion timing, with emergence tending to move from the scotophase towards the photophase as temperatures decreased. This trend is consistent with the daytime eclosion observed under conditions of conflicting photoperiodic and thermal cues, where the mean temperature was 18.5 °C.

Mechanistically, temperature can influence circadian timing at multiple levels, from post-transcriptional regulation of core clock components to circuit-level modulation of light responses. In *Drosophila*, thermosensitive splicing of a 3′-terminal intron in *period* contributes to temperature-dependent adjustments in activity timing (Chen et al. 2007), and temperature-sensitive neurons can gate light responsiveness in circadian circuits (Tian et al. 2025). While the molecular and neural substrates of photoperiodic-thermal integration in *L. sticticalis* remain unknown, the pronounced plasticity and cue-dependent structure of its eclosion rhythms provide an experimentally tractable phenotype for dissecting how light and temperature signals are integrated to shape developmental timing. Together, our results advance understanding of how biological clocks integrate light and temperature cues to regulate developmental timing in a migratory pest, with implications for phenological interpretation under changing environments and for informing the timing of pest forecasting and management efforts.

## Author contributions

FC, XJ and GW conceived and designed research. CL, ZX, HQ and JZ conducted experiments. FC and GW contributed new reagents and analytical tools. CL, ZX, GW, and FC analyzed data. CL, AC, and GW wrote the manuscript. FC, XJ, AC and GW revised and polished the manuscript. All authors read and approved the manuscript.

## Statements and Declarations

### Author contributions

Fajun Chen, Xingfu Jiang and Guijun Wan conceived and designed research. Changning Lv, Zun Xu, Hui Qiu and Jingxian Zhou conducted experiments. Fajun Chen and Guijun Wan contributed new reagents and analytical tools. Changning Lv, Zun Xu, Guijun Wan, and Fajun Chen analyzed data. Changning Lv, Abhishek Chatterjee, and Guijun Wan wrote the manuscript. Fajun Chen, Xingfu Jiang, Abhishek Chatterjee and Guijun Wan revised and polished the manuscript. All authors read and approved the final manuscript.

### Data availability

The datasets generated during the current study are available from the corresponding author on reasonable request.

### Ethics approval

Not applicable.

